# Whole genome bisulfite sequencing dataset of mycelium and spore of chalkbrood disease pathogen *Ascosphaera apis*

**DOI:** 10.1101/2020.03.23.002980

**Authors:** Yu Du, Haibin Jiang, Huazhi Chen, Jie Wang, Yuanchan Fan, Xiaoxue Fan, Cuiling Xiong, Yanzhen Zheng, Dafu Chen, Rui Guo

## Abstract

Chalkbrood, a widespread fungal disease of bee larvae, is caused by the fungus *Ascosphaera apis*. In this article, mecylia and spores of *A. apis* were respectively collected followed by DNA isolation, bisulfite conversion, cDNA library construction and next-generation sequencing. Using whole genome bisulfite sequencing (WGBS), 69,844,360 and 60,570,990 raw reads were yielded from Aam and Aas, and after quality control, 9,982,386,951 and 8,825,601,434 clean reads were obtained, respectively. In addition, 67,685,866 and 58,886,072 clean reads were mapped to the reference genome of *A. apis*, including 37,643,592 and 31,568,442 unique mapped clean reads, and 49,686 and 13,348 multiple mapped clean reads. Furthermore, after bisulfite treatment, the conversion ratio of clean reads from Aam and Aas were 99.38% and 99.51%, respectively. The WGBS data ducumented here contributes to genome-wide identification of 5mC methylation sites in *A. apis* and comparison of methylomes between mycelium and spore.

**Value of the data:**

- This dataset can be used for genome-wide identification of 5mC methylation sites in *A. apis*.
- The accessible data could be used to systematically compare methylomes between mycelium and spore of *A. apis*.
- Current data provides a useful resource for further study on DNA methylation-mediated mechanism underlying mycelium growth, spore germination and sexual reproduction of mycelium with the opposite sex.

## 1. Data description

The shared data were gained from whole genome bisulfite sequencing (WGBS) of *A. apis* mycelium and spore. As **Figure 1** presented, the ratio of G and C among total bases derived from WGBS were apparently much lower than that of A and T. Totally, 69,844,360 and 60,570,990 raw reads were yielded from Aam and Aas, amounting to approximately 10.48 Gb and 9.09 Gb, respectively (**Table 1**). After quality control, 9,982,386,951 and 8,825,601,434 clean reads were respectively obtained, amounting to about 9.98 Gb and 8.83 Gb (**Table 1**). In addition, the GC contents of clean reads from Aam and Aas were 27.52% and 23.37%, respectively (**Table 1**). As **Table 2** shown, 67,685,866 and 58,886,072 clean reads were mapped to the *A. apis* genome, including 37,643,592 and 31,568,442 unique mapped clean reads, and 49,686 and 13,348 multiple mapped clean reads. Moreover, after bisulfite treatment, the conversion ratio of clean reads from Aam and Aas were 99.38% and 99.51%, respectively.

**Figure 1.**
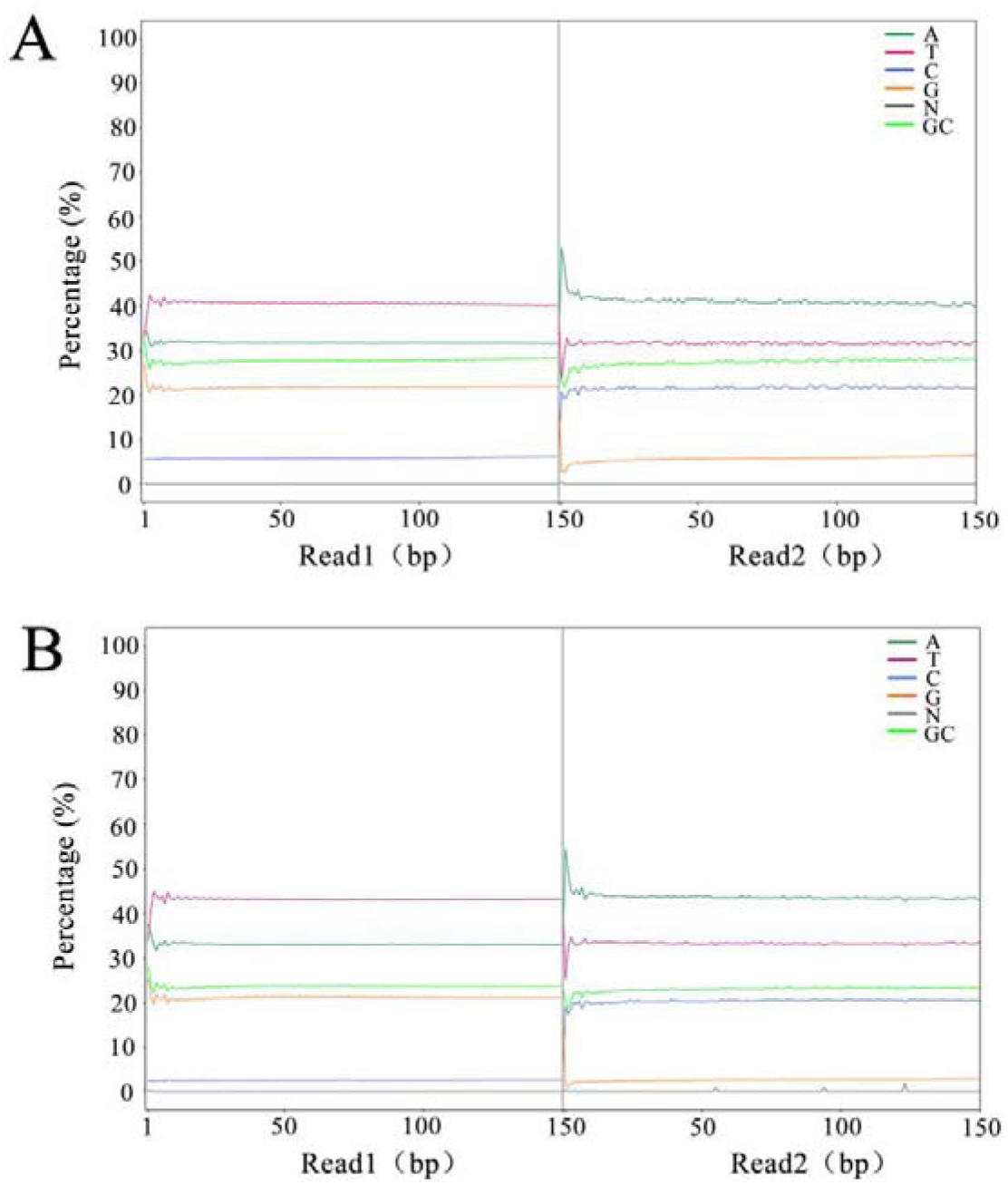
Ratio of four different bases from *A. apis* mycelium (A) and spore (B) after bisulfite conversion. The *x* axis indicates the position of reads, and the *y* axis indicates the ratio of a single base among total bases.

**Table 1.** Summary of WGBS of *A. apis* mycelium and spore.

| Sample | Raw reads | Clean reads | Percentage of clean reads | Total bases of raw reads | Total bases of clean reads | GC content |
| --- | --- | --- | --- | --- | --- | --- |
| Aam | 69,844,360 | 67,685,866 | 96.91% | 10,476,654,000 | 9,982,386,951 | 27.52% |
| Aas | 60,570,990 | 58,886,072 | 97.22% | 9,085,648,500 | 8,825,601,434 | 23.37% |

**Table 2.** Summary of mapping of clean reads to the reference genome of *A. apis*.

| Sample | Clean reads | Unique mapped clean reads | Multiple mapped clean reads | Mapping ratio |
| --- | --- | --- | --- | --- |
| Aam | 67,685,866 | 37,643,592 | 49,686 | 55.69% |
| Aas | 58,886,072 | 31,568,442 | 13,348 | 53.63% |

## 2. Experimental Design, Materials, and Methods

### 2.1. Sample preparation

*A. apis* was previously isolated from a fresh chalkbrood mummy of *Apis mellifera ligustica* larva [1] and kept at the Honeybee Protection Laboratory of the College of Animal Sciences (College of Bee Science) in Fujian Agriculture and Forestry University. Based on previously described approach [2] with several minor modifications [3], fungal mycelium sample and spore sample were respectively purified and immediately frozen in liquid nitrogen followed by storage at −80 °C until WGBS.

### 2.2. Bisulfite conversion and library preparation

Genomic DNA was respectively extracted from Aam and Aas using a Universal Genomic DNA Extraction Kit (TaKaRa, Tokyo, Japan) following the manufacturer’s protocol. DNA was bisulfite treated using a Zymo Research EZ DNA methylaiton-Glod Kits (Zymo Research, Irvine, CA, USA). DNA concentration and integrity were examined by a NanoDrop 2000 spectrophotometer (Thermo Fisher Scientific, Waltham, MA, USA) and agarose gel electrophoresis, respectively. Next, the libraries were constructed using TruSeq DNA Methylation Kit (Illumina, San Diego, CA, USA) according to the manufacturer’s instructions.

### 2.3. Deep sequencing and data processing

The libraries were sequenced on an Illumina HiSeq X Ten platform by OE Biotech Co., Ltd. (Shanghai, China) and 150 bp paired-end reads were generated. Clean reads were processed using fastp [4], which were obtained for downstream analyses by removing reads containing adapter, reads containing ploy-N and low quality reads from raw reads. Next, the clean reads were aligned to the *A. apis* genome (assembly AAP 1.0) using the WGBS mapping program (BSMAP, v2.74) [5]. Then, mapping ratio and bisulfite conversion rates were calculated for each sample.

## Acknowledgments

This research was financially supported by the Earmarked Fund for China Agriculture Research System (No. CARS-44-KXJ7), the Science and Technology Planning Project of Fujian Province (No. 2018J05042), the Teaching and Scientific Research Fund of Education Department of Fujian Province (No. JAT170158), the Outstanding Scientific Research Manpower Fund of Fujian Agriculture and Forestry University (No. xjq201814), and the Scientific and Technical Innovation Fund of Fujian Agriculture and Forestry University (No. CXZX2017342, No. CXZX2017343).

## Reference

[1] R. Guo, D.F. Chen, C.L. Xiong, C.S. Hou, Y.Z. Zheng, Z.M. Fu, Q.Y. Diao, L. Zhang, H.Q. Wang, Z.X. Hou, W.D. Li, D. Kumar, Q. Liang. Identification of long non-coding RNAs in the chalkbrood disease pathogen *Ascospheara apis*. J. Invertebr. Pathol. (2018), 156, 1–5, doi:http://doi.org/10.1016/j.jip.2018.06.001.

[2] A.B. Jensen, K. Aronstein, J.M. Flores, S. Vojvodic, M.A. Palacio, M. Spivak. Standard methods for fungal brood disease research. J. Apic. Res. (2013), 52(1), doi: http://doi.org/10.3896/IBRA.1.52.1.13.

[3] Y. Du, H.Z. Chen, J. Wang, Z.W. Zhu, C.L. Xiong, Y.Z. Zheng, D.F. Chen, R. Guo. Nanopore long-read transcriptome data of fungal pathogen of chalkbrood disease, *Ascosphaera apis*. bioRxiv (2020), doi: https://doi.org/10.1101/2020.03.12.989863.

[4] S. Chen, Y. Zhou, Y. Chen, J. Gu. fastp: an ultra-fast all-in-one FASTQ preprocessor. Bioinformatics, 2018, 34(17): i884–i890, doi: http://doi.org/10.1093/bioinformatics/bty560.

[5] Y. Xi, W. Li. BSMAP: whole genome bisulfite sequence MAPping program. BMC Bioinf. 2009, 10, 232, doi: http://doi.org/10.1186/1471-2105-10-232.

